# Capn5 expression in the healthy and regenerating zebrafish retina

**DOI:** 10.1101/274423

**Authors:** Cagney E. Coomer, Ann C. Morris

## Abstract

**Purpose:** Autosomal dominant neovascular inflammatory vitreoretinopathy (ADNIV) is a devastating inherited autoimmune disease of the eye that displays features commonly seen in other eye diseases, such as retinitis pigmentosa and diabetic retinopathy. ADNIV is caused by a gain of function mutation in Calpain-5 (*CAPN5*), a calcium dependent cysteine protease. Very little is known about the normal function of Capn5 in the adult retina, and there are conflicting results regarding its role during mammalian embryonic development. The zebrafish (Danio rerio) is an excellent animal model for studying vertebrate development and tissue regeneration, and represents a novel model to explore the function of Capn5 in the eye.

**Methods:** We characterized the expression of Capn5 in the developing zebrafish central nervous system (CNS) and retina, in the adult zebrafish retina, and in response to photoreceptor degeneration and regeneration using whole mount in situ hybridization, fluorescent in situ hybridization (FISH), and immunohistochemistry.

**Results:** In zebrafish, *capn5* is strongly expressed in the developing embryonic brain, early optic vesicles, and in newly differentiated retinal photoreceptors. We found that expression of *capn5* co-localized with cone specific markers in the adult zebrafish retina. We observed an increase in expression of Capn5 in a zebrafish model of chronic rod photoreceptor degeneration and regeneration. Acute light damage to the zebrafish retina, was accompanied by an increase in expression of Capn5 in the surviving cones and in a subset of Müller glia.

**Conclusions:** These studies suggest that Capn5 may play a role in CNS development, photoreceptor maintenance, and photoreceptor regeneration.

## Introduction

Calpains are a family of calcium-dependent, non-lysosomal cysteine proteases that include at least 15 family members in humans.^1-2^ Calpains share similarity in their protease and calcium binding domains, and are unique to other proteases because they are localized to the cytosol or nucleus instead of the lysosome. Activated by influxes of calcium, calpains do not degrade their protein substrates, but rather modify the activity of their targets through proteolytic processing. Calpains have been implicated in numerous cellular processes, including cell death, signal transduction, intracellular signaling, and sex determination.^3^ Hyperactivation of calpains is associated with the pathogenesis of several diseases, including Alzheimer’s, Parkinson’s, cardiovascular, and autoimmune diseases, as well as neurodegeneration following traumatic brain injury (TBI).^4^,^5^ Calpains have also been implicated in many eye and retinal diseases, such as retinitis pigmentosa, retinal detachment, and glaucoma.^6,7,8,9^ Although calpains have been the subject of extensive research, their functions and substrates are not fully defined. Moreover, while the majority of studies have focused on the ubiquitously expressed classical calpains CAPN1 (μ-calpain) and CAPN2 (m-calpain), the functions of the non-classical calpains in health and disease are much less well understood.

Capn5 is grouped with the non-classical calpains because it contains a C2-like domain instead of a penta-EF hand domain (domain 4) at its C-terminus. Capn5 is considered the vertebrate homolog of *C. elegans TRA3,* which plays a role in sex determination and mediates a necrotic pathway in neurons. ^10,11^ Capn5 has been shown to be the second most abundantly expressed calpain in the mammalian central nervous system (CNS).^12^ Expression of Capn5 has also been demonstrated in the mammalian retina, where it is found in the outer plexiform layer (OPL) and outer nuclear layer (ONL), specifically the inner and outer segments and synapses of the rod and cone photoreceptors, some ganglion cells and the inner plexiform layer.^13^ Within cells, Capn5 has been shown to be associated with promyelocytic leukemia protein bodies (PML) in the nucleus, which have been implicated in cellular stress response, apoptosis, cellular senescence, and protein degradation.^12-14^

Mutations in *CAPN5* are associated with the devastating retinal degenerative disease autosomal dominant neovascular inflammatory vitreoretinopathy (ADNIV).^15,16,17^ ADNIV is a hereditary autoimmune disease of the eye that is characterized by abnormal retinal pigmentation, retinal neovascularization, photoreceptor degeneration, vitreous hemorrhage, intraocular fibrosis, and tractional retinal detachment. As the disease progresses it phenocopies more commonly known ocular diseases such as non-infectious uveitis, glaucoma, diabetic retinopathy and retinitis pigmentosa. ^15,18^ The three point mutations that have been identified in ADNIV patients (p.Arg243Leu, p.Leu244Pro, and p.Lys250Asn) are located in the calcium-sensitive domain 2 near the active site and are thought to cause the mislocalization of CAPN5 from the cell membrane to the cytosol.^16,17^ Thus, ADNIV is thought to result from gain-of-function mutations in *CAPN5* that lower its threshold for activation by calcium.^15,19^ However, the precise mechanism whereby mutant CAPN5 causes ADNIV is not well understood. Elucidating the role of CAPN5 in the retina could reveal the underlying pathogenetic mechanisms of ADNIV as well as other retinal degenerative diseases that display similarities to ADNIV.

The normal function of Capn5 during development and in the adult retina is not well understood. Previous studies using two different *Capn5* mutant mouse models yielded conflicting results. In one study, *Capn5* null mice (*Capn5 trn1Nde*) were born viable and fertile, but some mutant offspring were runted and died two months after birth.^20^ In another study, the *Capn5-/-* null mutation (*Capn5 trn1Dgen*) was pre-implantation embryonic lethal.^20^ These conflicting results highlight a need for additional animal models to study Capn5 function. Zebrafish are an ideal model for developmental studies due to their strong genetic homology to humans, completely sequenced genome, and rapid, transparent, external development. Furthermore, unlike mammals, the zebrafish is capable of regenerating all classes of retinal neurons in response to injury or genetic insult. This regenerative response relies on the retinal Müller glia, which are stimulated to dedifferentiate, re-enter the cell cycle, and generate retinal progenitor cells to replace the cells that have been lost.^21-22^

In this study, we examined the expression profile of *capn5* during embryonic development and in the adult retina of the zebrafish. The zebrafish has two orthologs of *Capn5* (*capn5a* and *capn5b*; http://zfin.org/) and we demonstrate that both are expressed in the CNS and the retina during zebrafish development. We also demonstrate that Capn5 is expressed in the adult zebrafish retina in a similar pattern to that described for the mammalian retina, however we find that expression of Capn5 is specific to cone, but not rod, photoreceptors. Finally, we show that Capn5 is upregulated in response to photoreceptor degeneration and is expressed in the Müller glia following acute light exposure, suggesting that Capn5 could be playing a role in the regenerative process to retinal damage.

## Results

### Developmental expression of zebrafish *capn5* orthologs

While some studies have examined the expression and function of Capn1 and Capn2 orthologs in zebrafish,^23^ the developmental expression pattern of Capn5 has not previously been reported. Through BLAST searches of zebrafish genome databases we identified two, full-length cDNA sequences with significant sequence similarity to human *CAPN5*. Due to an ancient genome duplication that occurred during the evolution of teleost fishes,^24-25^ zebrafish often possess two orthologs of single-copy mammalian genes. The two zebrafish orthologs of *CAPN5*, hereafter referred to as *capn5a* and *capn5b,* are located on zebrafish chromosomes 18 and 21, respectively. The predicted zebrafish Capn5a protein is 68% identical and 81% similar to human CAPN5, whereas zebrafish Capn5b is 71% identical and 83% similar. We identified unique regions of each cDNA sequence and designed PCR primers. We extracted RNA from embryos at selected developmental time points (4, 18, 24, 48, 72, and 120 hours post fertilization), and performed RT-PCR followed by agarose gel electrophoresis to detect expression of *capn5a* and *capn5b.* Expression of both genes was detectable starting at 4 hours post fertilization (hpf), indicating *capn5* is maternally deposited, and both *capn5a* and *capn5b* were expressed at every subsequent developmental timepoint tested (Figure 1A). To quantify the expression of *capn5a* and *capn5b*, we performed quantitative real-time RT-PCR (qPCR). For both *capn5a* and *capn5b*, we observed a gradual increase in expression as development progressed (Figure 1B). With the exception of 4 hpf, expression of *capn5a* was consistently higher than *capn5b* (Figure 1B). From these data, we conclude that *capn5a* and *capn5b* are expressed during zebrafish embryonic development and that *capn5a* is expressed more strongly during development than *capn5b*.

**Figure 1:**
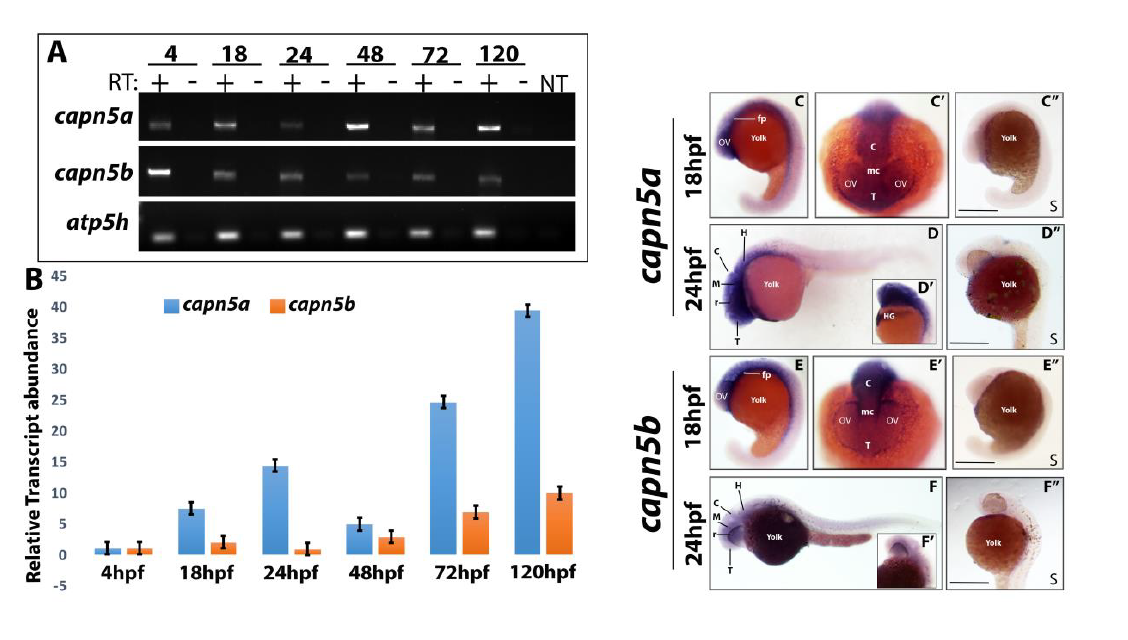
*capn5a* and *capn5b* are expressed in the developing brain of zebrafish. A) RT-PCR for *capn5a* and *capn5b* expression during development (24, 48, 72, and 120 hpf). *atp5h* expression is shown as housekeeping gene control. RT, reverse transcriptase; NT, no template. B) qPCR representation of fold-change in mRNA expression relative to 24 hpf. C-C") Whole-mount in situ hybridization (WISH) of *capn5a* at 18hpf. Strong *capn5a* expression was observed in the optic vesicles and brain. WISH with a control sense probe is shown C". D-D") WISH for *capn5a* at 24hpf. Strong expression of *capn5a* was detected in the brain and hatching gland. Control sense probe is shown D" E-E") WISH for *capn5b* at 18hpf. Expression was observed in parts of the developing brain but not the optic vesicles. Sense probe shown in E". F-F") WISH for *capn5b* at 24 hpf. Modest expression of *capn5b* was observed in the brain. (OV, optic vesicle; fp, floor plate; T, Telencephalon; mc, mesencephalon; c, cerebellum; HG, hatching gland; H, hindbrain; r, retina. Scale bars = 50μm.)

In the adult mouse, *Capn5* expression has been identified in many tissues including the brain, eye, uterus, and prostate;^12,26-29^ however, less is known about the expression pattern of *Capn5* during vertebrate embryonic development. To address this question, we performed whole-mount in situ hybridization (WISH) with unique probes for *capn5a* and *capn5b* at selected developmental timepoints in the zebrafish. At 18 hpf, *capn5a* expression was detected in the optic vesicles, developing diencephalon, mesencephalon and hindbrain (Figure 1C-C’). *Capn5b* expression was present in the diencephalon, mesencephalon, and hindbrain, but was not detected in the optic vesicles at 18 hpf (Figure 1E). At 24 hpf, expression of *capn5a* and *capn5b* is was observed in the developing zebrafish brain, more specifically the tectum, hindbrain, cerebellum and floor-plate. Expression of *capn5a* was also observed in the hatching gland (Figure 1D, 1F). We did not detect expression of *capn5a* or *capn5b* in the developing eye at 24, 36 or 48 hpf (data not shown). We conclude that *capn5a/b* are expressed in the developing zebrafish brain and in the optic vesicle prior to its invagination to form the bi-layered optic cup.

### *Capn5a* and *capn5b* are expressed in differentiated larval photoreceptors

To further investigate the expression of *capn5a* and *capn5b* in the zebrafish retina, we used fluorescent in situ hybridization (FISH) with *capn5a/b* probes on cryosections of zebrafish larvae at 120 hpf (5 days post fertilization, dpf). At 5 dpf, all of the retinal cell types have differentiated and zebrafish larvae display visually evoked behaviors. At this timepoint expression of both *capn5a* and *capn5b* was detected across the photoreceptor cell layer, in the region of the photoreceptor inner segments (Figure 2A-D’). In a previous study, Schaefer et al. demonstrated CAPN5 antibody localization to photoreceptor inner segments and synaptic terminals in adult mouse retinal sections.^13^ We used the same CAPN5 antibody for immunohistochemistry (IHC) on 5 dpf zebrafish retinal sections. This antibody should detect both Capn5a and Capn5b proteins. Similar to the previous study in mouse retina, we detected Capn5a/b protein expression in the photoreceptor inner segments and the outer plexiform layer (OPL) where the photoreceptor synaptic terminals are located. (Figure 2E-F’). We detected a similar pattern of expression of *capn5a/b* across the photoreceptor cell layer of the retina as early as 3 dpf (when the cone photoreceptors have largely finished differentiating), but not prior to this time point (data not shown). Therefore, we conclude that Capn5a/b is expressed in differentiated photoreceptors in the zebrafish retina. Furthermore, the lack of expression of *capn5a/b* in the retina prior to 3 dpf suggests that Capn5 plays a role in photoreceptor cell maintenance rather than development or specification.

**Figure 2:**
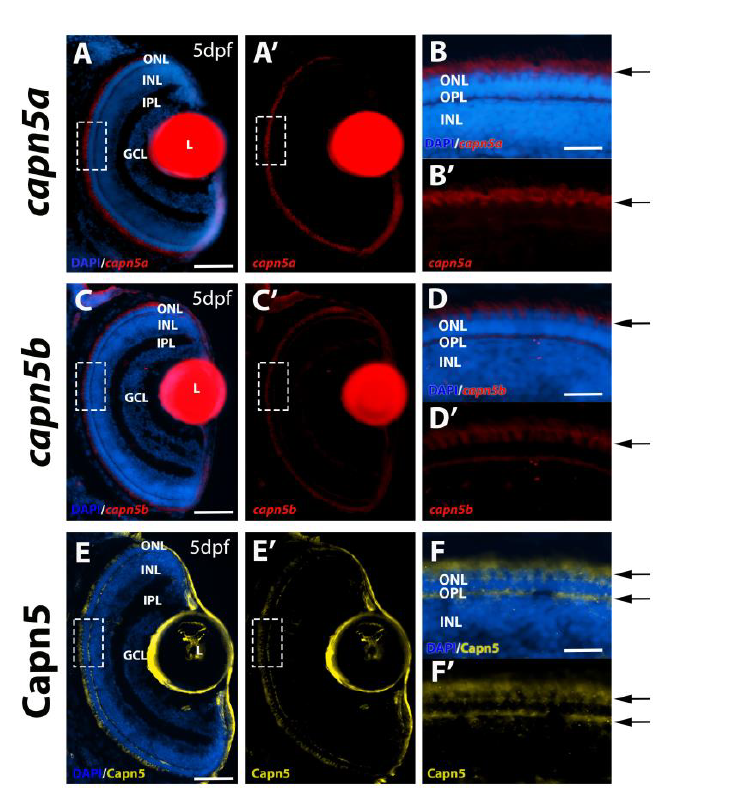
Capn5 is expressed in differentiated photoreceptors of the larval zebrafish retina. A-B’) Fluorescent in situ hybridization (FISH) showing expression of *capn5a* in the zebrafish retina at 5 dpf. Expression was detected in the outer nuclear layer (arrows). B-B’ show an enlarged image of the boxed area in A-A’. C-D’) FISH for expression of *capn5b* in the 5 dpf zebrafish retina. Expression was also detected in the outer nuclear layer (arrows). D-D’ show an enlarged image of the boxed area in C-C’. E-F’) Immunohistochemistry (IHC) showing expression of Capn5 in the outer plexiform layer and inner segments of the zebrafish photoreceptors at 5 dpf (arrows). F-F’ show an enlarged image of the boxed region in E-E’. L, lens (The red fluorescence in the Lens is non-specific autofluorescence); INL, inner nuclear layer; OPL, outer plexiform layer; GCL, ganglion cell layer; ONL, outer nuclear layer. Scale bars = 50 μm in A,C and E and 100 μm in B,D and F.

### Capn5 expression is cone-specific in the adult zebrafish retina

To further analyze the expression of *capn5a/b* in the adult zebrafish retina we performed FISH with *capn5a/b* probes on sections of adult wild type zebrafish retina. Expression of *capn5a* and *capn5b* was again observed in the photoreceptor inner segments (Figure 3). The rod and cone photoreceptors in the adult zebrafish retina are highly tiered and morphologically distinguishable, such that (moving from the vitreal to scleral direction), the round rod nuclei are located most proximal to the OPL, followed by the UV and blue cones, then the ellipsoid red/green double cone nuclei interspersed with rod inner segments, the red/green cone outer segments, and finally the rod outer segments located most proximal to the RPE.^30^ Interestingly, the FISH expression pattern for *capn5a/b* appeared to localize primarily to the cone, but not rod inner segments. To determine whether this was the case, we used the TαC-GFP transgenic line, in which GFP is expressed specifically in the cones,^31,32^ and performed FISH for *capn5a* in combination with IHC for GFP. We observed a perfect co-localization of GFP and *capn5a* expression (Figure 3D-D’’). We then performed the same experiment using the XOPS:GFP transgenic line, which expresses GFP specifically in the rods.^30^ We did not observe any co-localization of the rod GFP signal and *capn5a* expression (Figure 3E-E’’), which we confirmed using confocal microscopy (Figure 3F-F"). *Capn5b* displayed the same co-localization pattern as *capn5a* (not shown). We confirmed the expression of Capn5 protein in the adult retina using the CAPN5 antibody (Figure 3C-C”). Capn5 expression was observed in the cone inner segments, outer segments and in the synaptic terminals of the adult retina. Taken together, these results suggest that, similar to mammals, zebrafish Capn5 is expressed in retinal photoreceptors. However, zebrafish Capn5 expression appears to be cone-specific, which has not been reported for Capn5 in the mammalian retina.

**Figure 3:**
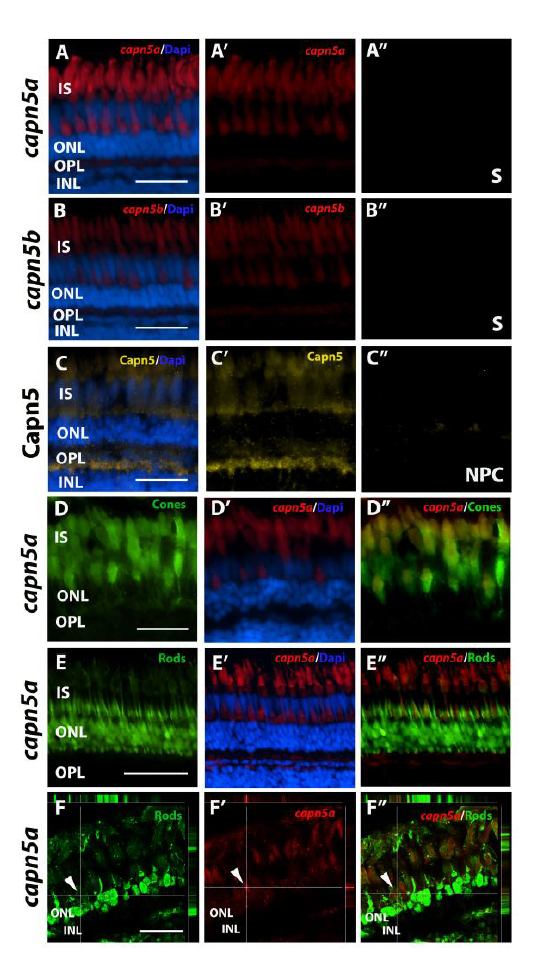
Capn5 is expressed specifically in the cone photoreceptors of the adult zebrafish retina. A-B’’) Fluorescent in situ hybridization (FISH) showing expression of *capn5a* and *capn5b* in the adult (WT) retina; expression was seen in the inner segments of the cones. A" and B" show sense probe controls. C-C’’) Immunohistochemistry (IHC) for Capn5 (both Capn5a and Capn5b). Expression was observed in the photoreceptor inner segments and in the outer plexiform layer. C" shows control image with no primary antibody. D-D’’) Co-localization of *capn5a* with a cone-specific marker. Combined FISH for *capn5a* and IHC for GFP on the T?C:GFP transgenic background demonstrates strong co-localization of *capn5a* expression with cone-specific GFP expression. E-E’’) Combined FISH for *capn5a* and IHC for GFP on the XOPS:GFP transgenic background demonstrates a lack of co-localization of *capn5a* expression with rod-specific GFP expression. F-F’’) Confocal microscopy imaging of combined FISH for *capn5a* and IHC for GFP on the XOPS:GFP transgenic background; the orthogonal view confirms there is no co-localization of rod specific GFP and *capn5a* expression. INL, inner nuclear layer; OPL, outer plexiform layer; ONL, outer nuclear layer; IS, inner segments; S, sense probe; NPC, no primary antibody control). Scale bars, 50 μm in E and 100 μm in A-D and F.

### *Capn5a* expression increases in response to rod photoreceptor degeneration

As described above, gain of function mutations in *CAPN5* have been associated with ADNIV, an autosomal dominant disease that eventually leads to the photoreceptor degeneration and retinal detachment.^12,16,17,19,33^ Moreover, other members of the calpain family have been implicated in apoptosis, necrosis, and photoreceptor degeneration.^6,7,34^ To determine whether Capn5 expression is altered in response to photoreceptor degeneration in zebrafish, we evaluated the expression of *capn5a* and *capn5b* in the XOPs:mCFP transgenic line, which displays continual degeneration and regeneration of rod (but not cone) photoreceptor cells.^35^ RT-PCR and qPCR were performed on mRNA prepared from dissected wild type (WT) and XOPS:mCFP retinas. In both experiments, we observed elevated expression of *capn5a* in the XOP:mCFP retinas compared to WT, whereas *capn5b* expression levels remained unchanged (Figure 4A-B). Next, we performed FISH on retinal cryosections from WT and XOPs:mCFP zebrafish with *capn5a/b* probes. We found that the expression patterns for both *capn5a* and *capn5b* were similar in the WT and the XOPs:mCFP retina; however, the signal for *capn5a* was stronger in XOPS:mCFP cones than in WT, whereas *capn5b* expression levels did not change (Figure 4C-H’). These results indicate that the cone-specific expression of *capn5a* is upregulated in response to rod photoreceptor degeneration in zebrafish.

**Figure 4:**
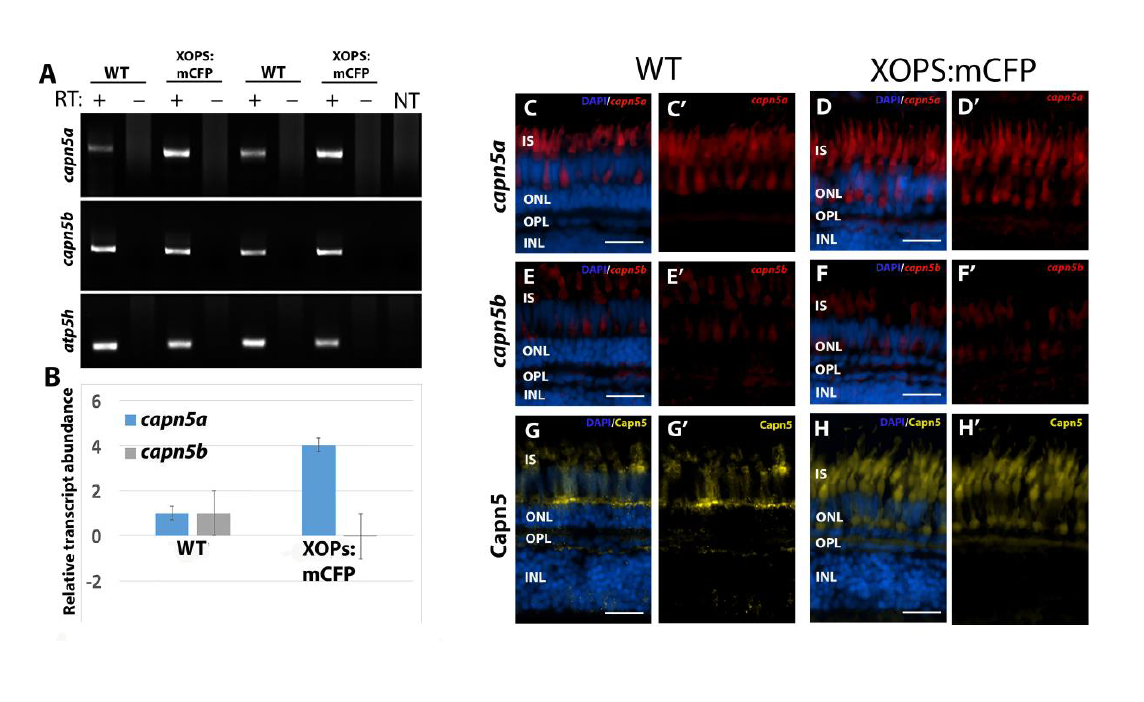
Expression of Capn5 is elevated in the adult XOPS:mCFP retina. A) RT-PCR for *capn5a* and *capn5b* expression in wild type (WT) and XOPS:mCFP retinas. *Capn5a* and *capn5b* are both expressed in the adult WT retina and *capn5a* levels are increased in the XOPS:mCFP retina. RT, reverse transcriptase; NT, no template control. B) qPCR for *capn5a* and *capn5b* in WT and XOPS:mCFP adult retina; a 4-fold increase of *capn5a* expression was observed in the XOPS:mCFP retina compared to WT. C-C’) Fluorescent in situ hybridization (FISH) showing expression of *capn5a* in the adult (WT) retina; expression was seen in the inner segments of the cones. D-D’) FISH showing expression of *capn5a* in the XOPS:mCFP retina; expression in the cone inner segments appears to be more intense than in WT. E-E’) FISH for *capn5b* in the adult WT retina; modest expression was detected in the inner segments. F-F’) FISH for *capn5b* in the XOPS:mCFP retina; expression levels appears similar to WT. G-G’) Immunohistochemistry (IHC) for Capn5 (a and b) in the WT adult retina; expression was detected in the outer plexiform layer (OPL) and in the cone inner segments (IS). H-H’) IHC for Capn5 in the XOPS:mCFP retina; increased expression was detected in the OPL and IS. INL, Inner nuclear layer; OPL, outer plexiform layer; ONL, outer nuclear layer; IS, photoreceptor inner segments. Scale bars, 100 μm.

### *Capn5* expression increases in response to acute light damage

Our results indicate that cone-specific expression of Capn5 is induced in the XOPS:mCFP transgenic line. However, in the XOPs:mCFP retina, rod photoreceptor degeneration does not result in any secondary degeneration of the cones.^35,36^ This led us to ask whether upregulation of Capn5 in cones serves a cell-autonomous protective function, or whether it plays a non-cell autonomous role in promoting rod degeneration or regeneration. To begin to address this question, we used an acute light damage approach, which allowed us to temporally separate photoreceptor degeneration and regeneration. We adopted the light damage protocol described by Vithelic and Hyde,^37^ in which dark adapted albino zebrafish are exposed to 20,000 lux of constant light for three days, followed by 7 days of recovery in normal lighting conditions. The acute light exposure causes almost total ablation of the rods and extensive damage to the cones in the dorsal retina, and less severe damage to rods and cones in the ventral retina.^38^ This is accompanied by de-differentiation and proliferation of a subset of the Müller glia, which produce retinal progenitor cells that migrate to the ONL to replace the degenerated photoreceptors. Zebrafish were collected prior to the start of the acute light damage, on the third day of light damage (LD), two days post-LD, and seven days post-LD, and the retinas were dissected and processed as described above for RT-PCR, qPCR, FISH and IHC.

RT-PCR and qPCR revealed a 12-fold upregulation of *capn5a* expression during photoreceptor degeneration (LD) followed by a return to WT levels by 7 days post-LD; we saw no change in *capn5b* expression during the entire light damage experiment (Figure 5A-B). To determine whether other calpains are also upregulated in response to retinal light damage, we analyzed the expression of *capn1* and *capn2* by RT-PCR (data not shown) and qPCR. We observed no significant increase in expression of *capn1a/b* or *capn2a/b* during light damage. *Capn1a/b* expression also did not change at either time point post-LD. However, we did observe a 3.5-fold increase in expression of *capn2a/b* at two and seven days post-LD (Supplemental Figure 1). Thus, the increase in expression observed during light damage is specific to *capn5a*.

**Figure 5:**
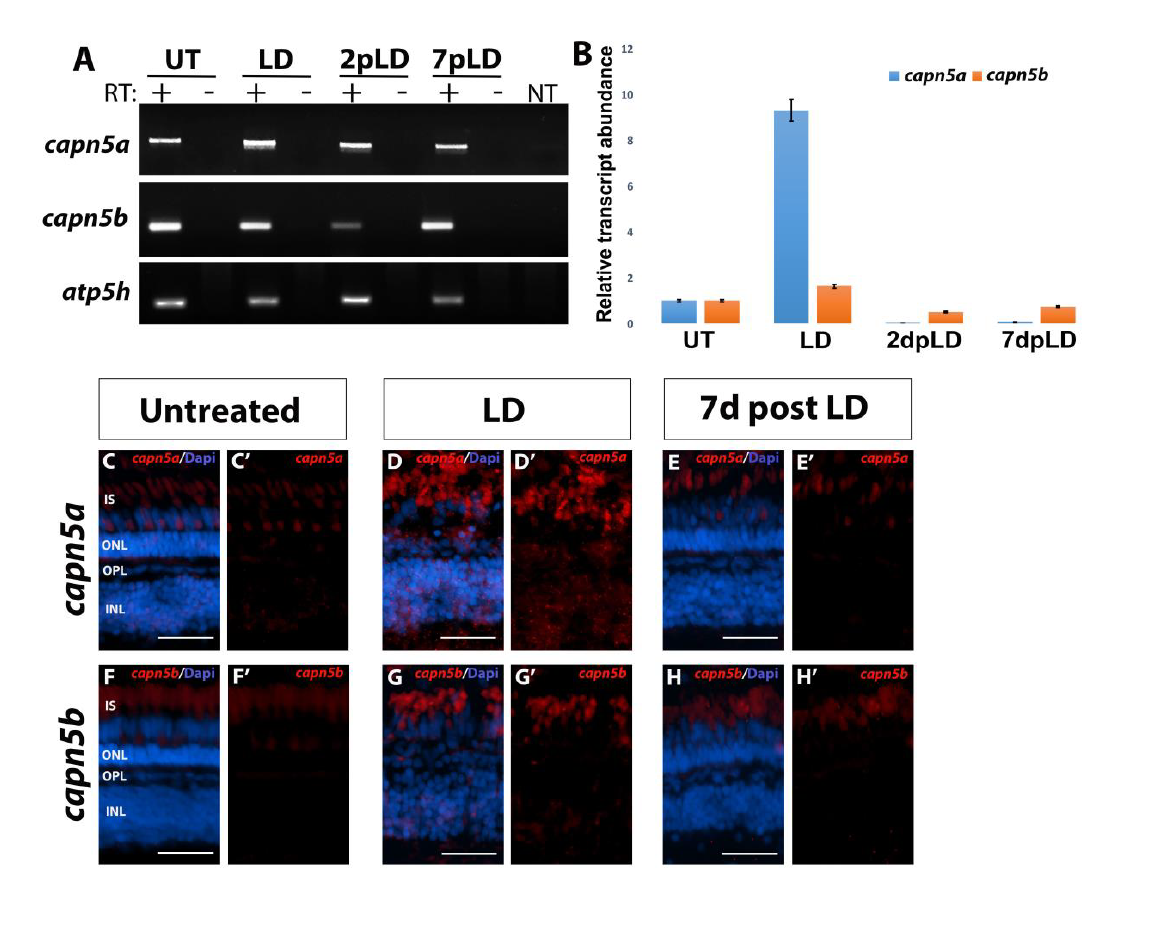
Capn5a expression is induced in response to acute light damage. A) RT-PCR of *capn5a* and *capn5b* expression during and following acute light damage. A significant increase in expression of *capn5a* was observed after 3 days of LD (LD) which returned to WT levels by 2dpLD. *Capn5b* expression did not change during LD. RT, reverse transcriptase; NT, no template control. B) qPCR qPCR for *capn5a* and *capn5b* during and following light damage. A 12-fold increase in *capn5a* expression was observed in the LD retina compared to untreated retina, followed by a significant decrease in expression post LD. *Capn5b* expression did not change during or after LD. C-E’) Fluorescent in situ hybridization (FISH) for *capn5a* in untreated retina (UT), during light damage (LD), and post light damae (7dpLD). Whereas *capn5a* expression in the UT retina was confined to the outer plexiform layer (OPL) and cone inner segments (IS), *capn5a* expression in the LD retina was seen in the inner nuclear layer (INL) as well as the OPL and IS. F-H’) FISH for *capn5b* in the UT, LD, 7dpLD retina. The *capn5b* expression pattern did not change during or after LD. INL, inner nuclear layer; OPL, outer plexiform layer; ONL, outer nuclear layer; IS, photoreceptor inner segments. Scale bars, 50 μm.

We performed FISH to characterize the expression pattern of *capn5a/b* in the retina during and following light damage. After three days of LD, the ONL was severely disrupted, with a significant reduction in the number of rod and cone nuclei. This was accompanied by novel expression of *capn5a* in the inner nuclear layer (INL) and increased expression in the remaining cone photoreceptors (Figure 5D-D’). By 7 days post-LD, the number of rod and cone nuclei had increased and the ONL appeared more organized. At this time point, expression began to return to the ONL only (data not shown) and by 7dpLD expression of *capn5a* was no longer observed in the INL and the photoreceptor expression resembled the pre-LD pattern (Figure 5E-E’). Expression of *capn5b* was restricted to the photoreceptor layer throughout the light damage experiment (Figure 5F-H’). Taken together, we conclude that *capn5a* expression increases during photoreceptor degeneration in response to acute light damage, with the novel INL expression unique to *capn5a*. Meanwhile, *capn5b* expression does not change in response to light-induced photoreceptor degeneration and regeneration.

### Expression level of Capn5 correlates with the extent of photoreceptor degeneration

As mentioned above, acute light damage results in greater photoreceptor degeneration in the dorsal versus the ventral retina.^38^ This regional difference allowed us to determine whether the extent of Capn5 expression is correlated with the level of photoreceptor damage. Using the Capn5 antibody (which detects both Capn5a and Capn5b), we compared Capn5 protein expression in the dorsal and ventral retina by IHC prior to LD, at 3 days of LD, and at 7 days post-LD. IHC for cone and rod specific markers was used to assess the amount of damage induced in the retina. In untreated retinas, as we observed previously, Capn5 was expressed in the cone photoreceptor inner segments and synaptic terminals (Figure 6A-F’). Capn5 expression appeared to be stronger in the dorsal retina than in the ventral retina (Figure 6E-F’). After three days of LD rod photoreceptors were completely ablated and cone photoreceptors were reduced and severely truncated in the dorsal retina (Figure 6G-I’). In the dorsal region, Capn5 expression was upregulated not only in the photoreceptor cell layer, but also in a subset of cells in the INL with a morphology suggestive of Müller glia (Figure 6K-K’). In contrast, in the ventral retina the rods were reduced and severely distorted, but the cones were much less damaged. In this region, Capn5 expression remained strong in the photoreceptor layer but was much weaker in the INL (Figure 6H-L’)

**Figure 6:**
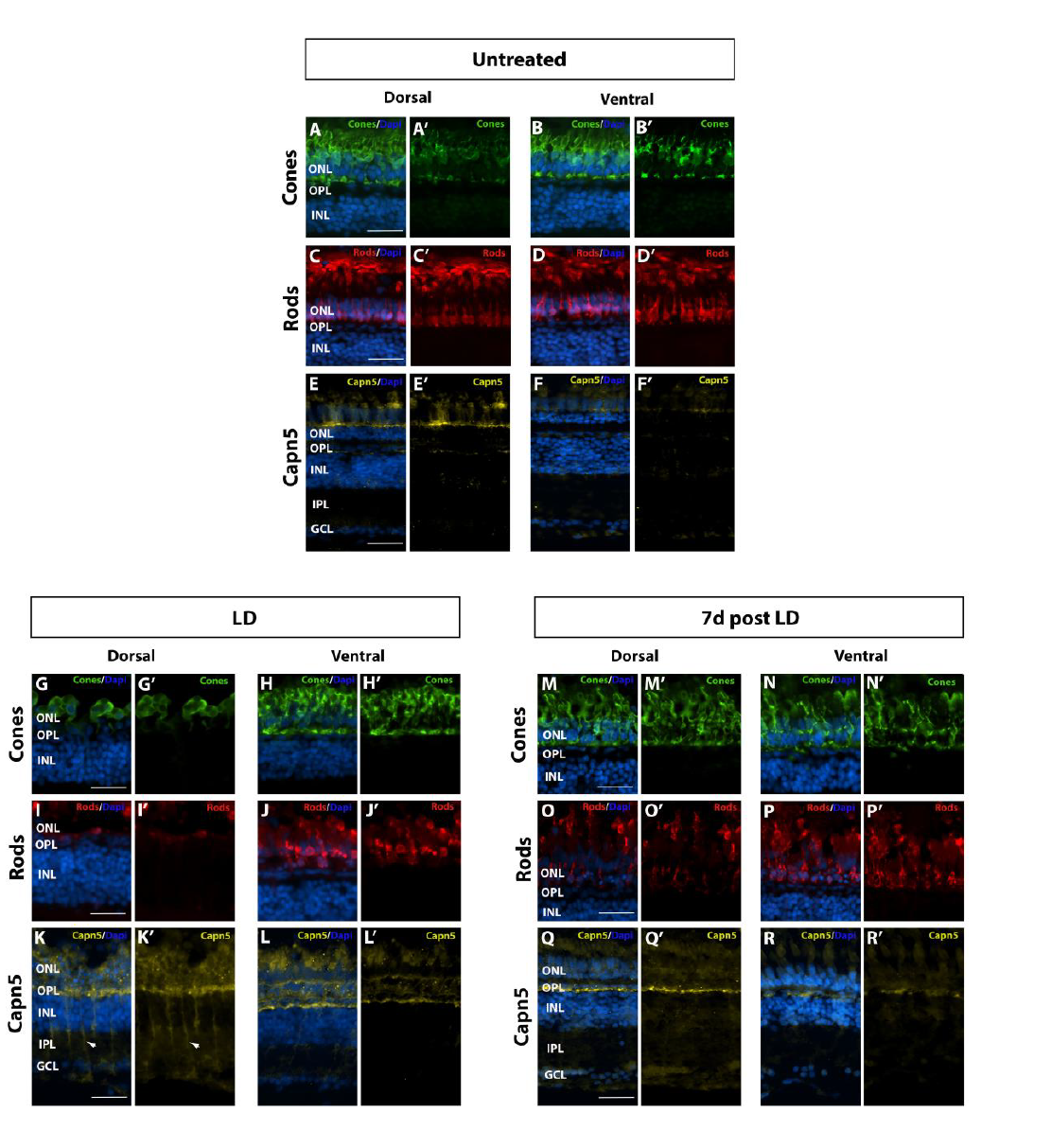
Increase in Capn5 expression correlates with magnitude of retinal damage. A-B’) Dorsal and ventral expression of cone specific marker Zpr-1 in undamaged (UT) retina. C-D’) Dorsal and ventral expression of rod specific marker 4C12 in the UT retina. E-F’) IHC for dorsal and ventral Capn5 expression in the UT retina; Capn5 expression is stronger in the dorsal retina (E-E’) compared to the ventral retina (F-F’). G-H’) Dorsal and ventral cone damage after acute light exposure. There is a significant decrease in the amount of cone photoreceptors in the dorsal retina compared to the ventral retina. (I-J’) Dorsal and ventral rod damage after acute light exposure. Rods are almost totally ablated in the dorsal retina, with more moderate damage observed in the ventral retina. (K-L’) IHC for Capn5 expression in the LD retina. Capn5 is strongly upregulated in the inner nuclear layer and surviving cones in the dorsal retina, and more modestly upregulated in the ventral retina. (M-N’) Dorsal and ventral cones have mostly recovered by 7 days post LD. O-P’) At 7 days post LD rods are regenerating, with an more rods observed in the ventral compared to the dorsal retina. Q-R’) IHC for Capn5 expression in the 7dpLD retina. Expression of Capn5 is similar to that of the undamaged retina. GCL, ganglion cell layer; INL, inner nuclear layer; OPL, outer plexiform layer; ONL, outer nuclear layer; Scale bars = 50 μm.

Finally, at 7 days post-LD, both rods and cones reappeared in the dorsal retina, and were very abundant in the ventral retina (Figure 6M-P’). This was accompanied by a disappearance of Capn5 expression from the INL and a decrease in expression in the photoreceptor layer in the dorsal retina (Figure 6Q-Q’), as well as a return to pre-LD expression levels in the ventral retina (Figure 6R-R’).

We conclude that photoreceptor degeneration caused by acute light damage induces Capn5 expression in proportion to the level of damage inflicted. Moreover, when photoreceptor degeneration is severe, Capn5 expression is upregulated in a subset of cells with a Müller glia morphology as well as in the ONL.

### Photoreceptor degeneration induces the expression of Capn5 in Müller glia

Finally, to confirm the identity of the Capn5-expressing cells in the INL of light damaged retinas, we performed co-localization experiments with the Capn5 antibody and the Zrf-1 antibody, which labels the processes of the Müller glia. In untreated retinas, we were unable to detect Capn5 expression in the INL and there was no co-localization of Capn5 expression with the Müller glia marker (Figure 7A’-C’). In contrast, after three days of LD, we observed an upregulation of Capn5 in the INL, and an increase in Zrf-1 staining indicative of the reactive gliosis that occurs in response to acute retinal damage.^39^ Moreover, we observed strong co-localization of the Capn5 signal with Zrf-1 (Figure 7D-F’). These results demonstrate that Capn5 is induced in Müller glia in response to photoreceptor degeneration.

**Figure 7:**
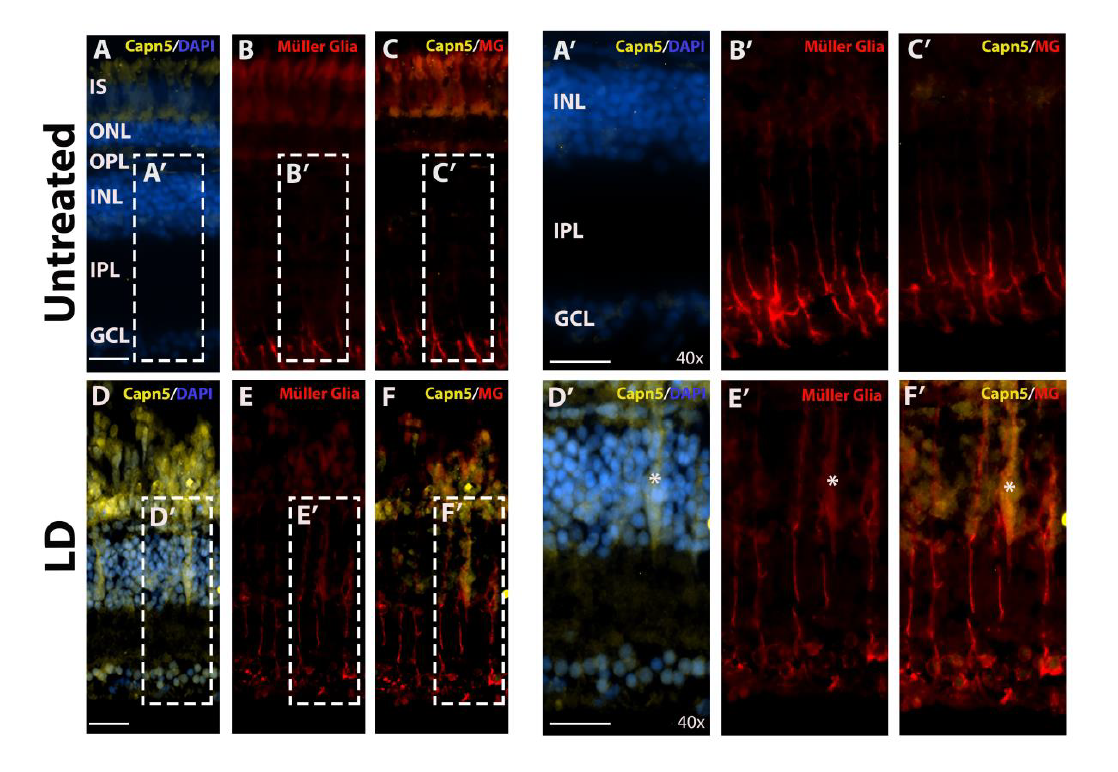
*capn5a* is expressed in the Müller glia in response to acute light damage. (A-C’) Immunohistochemistry (IHC) for Capn5 expression (A) and the Müller glial marker Zrf-1 (B) in the undamaged (UT) adult retina. No co-localization of Capn5 and Müller cells is observed. A’-C’ show an enlarged image of the boxed regions in A-C. D-F) IHC for Capn5 and Müller glia in the LD retina. Capn5 expressed in induced in the INL, where is co-localizes with reactive Müller glia (asterisk). D’-F’ show enlarged images of the boxed regions in E-F. GCL, Ganglion cell layer; IPL, inner plexiform layer; INL, inner nuclear layer; OPL, outer plexiform layer; ONL, outer nuclear layer; IS, photoreceptor inner segments. Scale bars, 100 μm in A-F and 500 μm in A’-F’.

### Discussion

Although mutations in CAPN5 are associated with a severe retinal degenerative disease, very few studies have described the developmental expression pattern of Capn5. In the mouse, developmental expression of Capn5 was suggested to be relatively low,^27^ with specific sites of expression noted in the developing thymus and the neurons of the sympathetic and dorsal root ganglia.^27,40^ Further in situ hyrbridization studies on embryonic tissue sections revealed *Capn5* expression in the developing mouse brain and in epithelial cells surrounding several tissues.^20,28^ In our study, we found that Capn5 is expressed in the developing brain at 18 hpf, and its expression increases throughout embryogenesis. We detected expression of Capn5 in the mesencephalon, telencephalon and optic vesicles at 18 hpf, and the hindbrain and cerebellum at later stages. This expression pattern supports a role for Capn5 during vertebrate brain development.

Cell death is a common and often essential process throughout embryonic development,^41^ and calpains have been implicated in mediating cell death pathways. Calpain 1 has been shown to cleave procaspase-12, activating caspase-12, and initiating cell death.^42^ In another study, calpain 2 was shown to mediate the cleavage of Atg5 switching the cell death pathway from autophagy to apoptosis.^4,43^ Given that calpains are thought to modify a wide variety of target substrates, we hypothesize that Capn5 plays a role in mediating cell death pathways in the nervous system during embryonic development.

To our knowledge, there are no published studies analyzing the expression of Capn5 during retinal development. Since mutations in Capn5 have been shown to cause the retinal degenerative disease ADNIV,^17^ elucidating the expression of Capn5 during retinal development is imperative. Our data reveal that in the zebrafish retinal expression of Capn5 is detectable in photoreceptors at 72 hpf, a time point at which the majority of cells have exited the cell cycle and differentiated. This indicates that Capn5 is not needed for the development or specification of photoreceptors, but rather that Capn5 is important for mature photoreceptor function or maintenance.

Previous studies have analyzed the expression of Capn5 in the adult human and rat retina, identifying protein expression in the outer nuclear layer, the outer plexiform layer, and the inner segments of the photoreceptors, as well as lower expression in some ganglion cells and the inner plexiform layer.^12^ Co-localization of Capn5 in the outer plexiform layer (OPL) with PSD95 expression, a marker for neural synaptic densities, coupled with the detection of Capn5 in the synaptic membrane by Western blot, indicated that Capn5 is expressed in the photoreceptor synapses. More specifically, expression was identified in both the outer segment and synaptic fractions of rod photoreceptors from mouse retina.^13^ In our study, we found that Capn5 is also expressed in mature photoreceptors in the wild type adult zebrafish retina, however its expression appears to be cone specific. We observed *capn5* mRNA expression in the inner segments of the cone photoreceptors, and analysis of protein expression by immunohistochemistry identified expression in the inner segments and the photoreceptor synaptic terminals, which is consistent with the mammalian expression pattern retina.^13^

Previous studies of Capn5 expression in the mouse and human retina did not report whether Capn5 expression was associated with a specific photoreceptor subtype. The zebrafish possesses a cone-rich retina with each of the four spectral subtypes having distinct morphologies, making them easily identifiable in tissue sections. We found, using both FISH and immunohistochemical analyses, that Capn5 is not expressed in the rod photoreceptors of the zebrafish. It will be interesting to determine whether mammalian Capn5 is also expressed primarily in cones, or whether there are species-specific differences in its localization.

Using a genetic model of rod-specific degeneration and regeneration,^35^ we found that the cone-specific expression of Capn5 increases in response to rod degeneration. This is intriguing because the cone photoreceptors in the XOPs-mCFP zebrafish do not degenerate secondary to the rod degeneration, as is typically observed in other photoreceptor degeneration and disease models.^35^ This raises the question: Is Capn5 playing an anti-apoptotic role in the cone photoreceptors and protecting them from the residual effects of the rod degeneration?

Using an acute light-damage paradigm, which results in almost total rod photoreceptor loss and some cone photoreceptor damage, we found that there is a large increase in Capn5 expression during the acute light damage phase that correlates with the amount of photoreceptor damage produced. In the zebrafish retina, the cones are much more resistant to the toxic effects of acute light exposure. Therefore, induction of Capn5 in cones in response to acute light damage could indicate a protective role for this protein during photoreceptor degeneration.

How do our data fit into the context of previously described functions of calpains? The role of calpains in regulating cell death pathways such as apoptosis, programmed cell death and necrosis has been extensively explored,^44^ with a focus on their function as either pro-or anti-apoptotic proteases. It has been shown that calpains play a pro-apoptotic role in the presence of a wide variety of stimuli. For example, in fibroblasts overexpression of Calpastatin, the endogenous calpain inhibitor, protected cells from okadaic acid induced apoptosis.^45^ Depletion of calpastatin in neutrophils promoted cell death during cycloheximide induced apoptosis.^46^ Evidence has also been presented that calpains play a pro-apoptotic role during cell death induced by hydrogen peroxide, ultra-violet light, and serum starvation through the PI3-kinase/Akt survival pathway.^1,26^ Overexpression of Capn2 in CHO cells resulted in sensitivity to ER stress induced cell death.^44^ In contrast, anti-apoptotic roles for calpains have also been demonstrated in the presence of some stimuli. For example, Capn1 cleaves p53 which protects the cell from DNA damage induced apoptosis.^47,48^ Given that no Capn5-specific substrates have been identified yet, it is possible that Capn5 plays an apoptotic or anti-apoptotic role, depending on the context. As mentioned above, our data indicate a possible protective role for Capn5 in the cones in response to rod photoreceptor degeneration. Future studies could test this hypothesis by creating a cone-specific Capn5 knockout and inducing photoreceptor cell death using acute light damage. If loss of Capn5 in cone photoreceptors results in a lower light threshold to induce cone cell death and/or an increase in the amount of cone damage caused by acute light exposure, this would support a protective function of Capn5 in cones.

Finally, one of the most intriguing results from our study is that, in addition to upregulation of Capn5 expression in the cones in response to acute light damage, we also observed induction of Capn5 in a subset of Müller glia. It should be noted that this Müller glia expression was not observed in the wild type or XOPs-mCFP zebrafish models, and has not been observed in the wild type mammalian retina. In teleosts, Müller glia are the source of retinal stem and progenitor cells for injury-induced regeneration, and they also phagocytose cell debris to clean up the retina in the initial response to damage.^49,50^ Therefore, the upregulation of Capn5 in Müller glia in response to acute light damage suggests a potential role for this calpain in photoreceptor regeneration as well.

In summary, this body of work lays the foundation for understanding the physiological function of Capn5 in the developing retina and in response to photoreceptor degeneration. As one of the few calpains that is not tightly associated with its inhibitor, understanding the function of Capn5 provides an opportunity to investigate calpain protease function in the absence of endogenous inhibition. Further investigation has the potential to shed light on the importance of proteolytic proteases in degeneration and regeneration, and possibly unlock the underlying mechanism associated with ADNIV.

## Methods

### Zebrafish lines and maintenance

All zebrafish lines were bred, housed, and maintained at 28.5°C on a 14 hour light:10 hour dark cycle, except where indicated for the light damage experiments. The Tg(XRho:gap43-mCFP) q13 transgenic line (hereafter called XOPS:mCFP) has been previously described.^35,36,51^ The Tg(3.2TαC:EGFP) transgenic line (TαC:EGFP) has been previously described,^31^ and was generously provided by Susan Brockerhoff (University of Washington, Seattle, WA). The Tg(XlRho:EGFP) transgenic line (hereafter called XOPS:GFP) has been previously described,^30^ and was obtained from James Fadool (Florida State University, Tallahassee, FL). Zebrafish were bred, raised and maintained in accordance with established protocols for zebrafish husbandry.^52^ Embryos were anaesthetized with Ethyl 3-aminobenzoate methanesulfonate salt (MS-222, Tricaine, Sigma-Aldrich, St. Louis, MO) and adults were euthanized by rapid cooling as previously described.^53^ All animal procedures were carried out in accordance with guidelines established by the University of Kentucky Institutional Animal Care and Use Committee (IACUC) and the ARVO Statement for the Use of Animals in Ophthalmic and Vision Research.

### RNA extraction, RT-PCR and Real-time quantitative RT-PCR (qPCR)

Total RNA was extracted from whole embryos at selected developmental time points or from the dissected retinas of adult zebrafish using TRIzol reagent (Invitrogen, Grand Island, NY) according to the manufacturer’s protocol. The samples were treated with RNAse-free DNAse I (Roche, Indianapolis, IN) and purified using a chloroform/phenol extraction. The GoScript Reverse Transcriptase System (Promega, Madison, WI) was used to synthesize first strand cDNA from 1μg of the extracted RNA. PCR primers were designed to amplify unique regions of the *capn5a, capn5b,* and *atp5h* cDNAs (Eurofins Genomics; www.eurofinsgenomics.com; Table 1). Faststart Essential DNA Green Master mix (Roche) was used to perform qPCR on a Lightcycler 96 Real-Time PCR System (Roche). The relative transcript abundance was normalized to *atp5h* expression as the housekeeping gene control,^54^ and was calculated as fold-change relative to 4 hours post fertilization (hpf) for developmental expression, and fold-change relative to wild type, untreated adult fish (WT) for the XOPs:mCFP and light damage experiments. RT-PCR and qPCR experiments were performed with three biological replicates and three technical replicates. RT-PCR was performed on a Mastercycler Pro thermocycler (Eppendorf, Westbury, NY). PCR products were visualized on a 1% agarose gel. The sequences for the primers used to produce the PCR products are listed in Table 1.

**Table 1:**
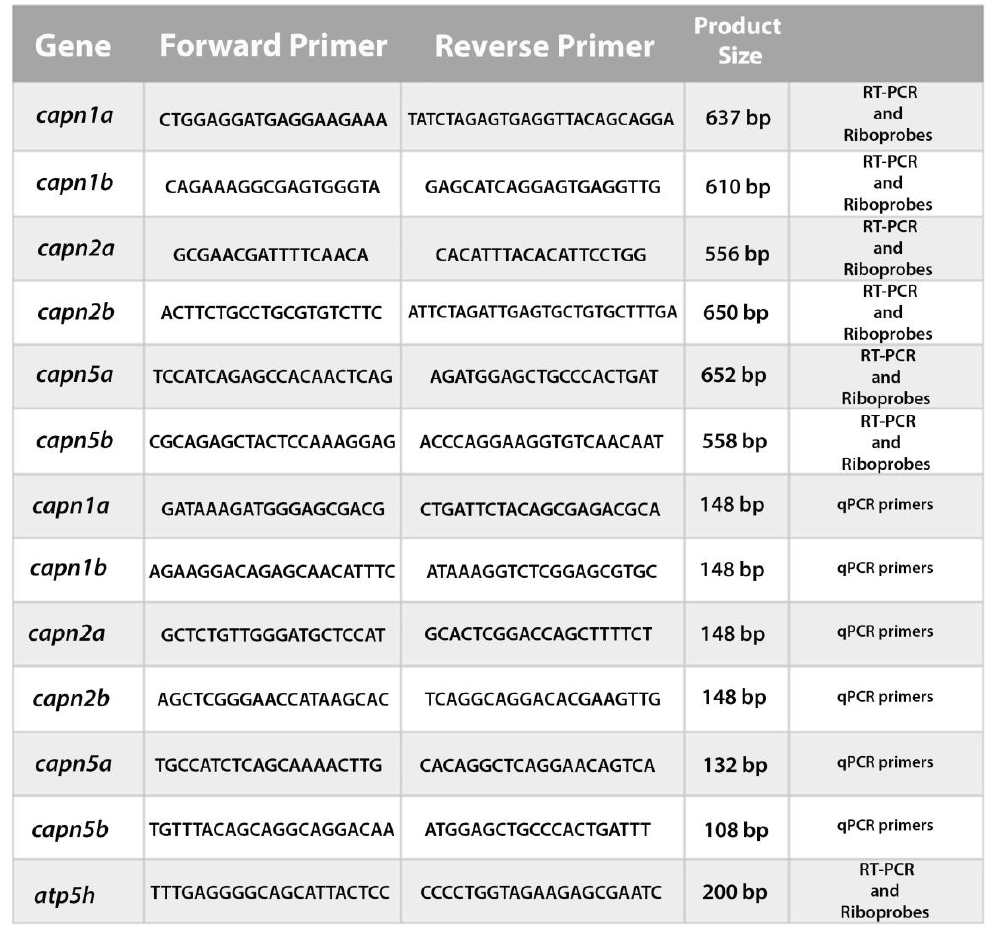
Primer sequences used for RT-PCR and qPCR. RT-PCR primers were also used to design WISH and FISH probes.

### Tissue sectioning

Whole embryos and adult retinas were collected as described above and fixed in 4% paraformaldehyde (PFA) at 4°C overnight. Fixed embryos or retinas were cryoprotected in 10% sucrose for a minimum of 8 hours, followed by 30% sucrose overnight at 4°C. Samples were placed into OTC medium (Ted Pella, Redding, CA) and frozen at −80°C for 2 hours. Ten micron-thick tissue sections were cut on a cryostat (Leica CM 1850, Leica Biosystems, Buffalo Grove, IL) and the sections were mounted on gelatin-coated or Superfrost Plus slides (VWR, Randor, PA) and air-dried overnight at room temperature.

### Riboprobe synthesis

PCR products from the unique regions of *capn5a* and *capn5b* were cloned into the pGEMT-easy vector (Promega, Madison, WI). Plasmids were linearized using either SpeI or SacII restriction enzymes (NEB, Ipswich, MA). Riboprobes were synthesized from the plasmids by in vitro transcription using either T7 or Sp6 polymerase and a digoxigenin (DIG) labeling kit (Roche Applied Science, Indianapolis, IN). The sequences for the primers used to produce the PCR products are listed in Table 1.

### Fluorescent in situ hybridization (FISH)

FISH was performed essentially as previously described.^55,56^ Briefly, embryos or adult retinas were fixed and sectioned as described above. Sections were post-fixed in 1% PFA and rehydrated in PBST. All solutions were prepared with diethyl pyrocarbonate (DEPC)-treated water. Hydrated sections were permeabilized for 10 minutes with 1μg/ml proteinase K, then acetylated in triethanolamine buffer containing 0.25% acetic anhydride (Sigma-Aldrich, Saint Louis, MO) and rinsed in DEPC treated water. Sections were incubated with DIG-labeled riboprobes (final concentration of 3 ng/μl) in hybridization buffer at 65°C in a sealed humidified chamber for at least 16 hours. After hybridization the sections were rinsed in 5x SSC followed by a 30 min 1xSSC/ 50% formamide wash. 1% H2O2 was used to quench the peroxidase activity and the sections were blocked using 0.5% PE blocking solution (Perkin Elmer Inc, Waltham, MA) for a minimum of 1 hour. Sections were incubated with anti-DIG-POD fab fragment (Roche, Indianapolis, IN) at 4°C overnight. The TSA plus Cy3 Kit (Perkin Elmer Inc, Waltham, MA) was used for probe detection. The sections were counterstained with 4’, 6-diamidino-2-phenylindole (DAPI, 1:10,000 dilution, Sigma-Aldrich), mounted in 60% glycerol, and imaged on an inverted fluorescent microscope (Nikon Eclipse Ti-U, Nikon instruments, Melville, NY) using a 20x or 40x objective or a Leica SP8 DLS confocal/digital light sheet system (Leica Biosystems, Nussloch, Germany) using a 40x or 60x objective. At least 3 retinas or 6 embryos and a minimum of 6 sections were analyzed for each timepoint and probe.

### Whole-mount in situ hybridization (WISH)

Embryos were manually dechorionated, collected at selected developmental time points (18, 24, 48 hpf, 72 hpf and 120 hpf) and fixed as described above. WISH was performed as previously described. ^55^ DIG-labeled riboprobes (3 ng/μl) were hybridized to the samples overnight at 60°C in hybridization buffer. After washing and blocking, samples were incubated overnight at 4°C with an anti-DIG-AP antibody (Roche, diluted 1:2000 in blocking solution). The next day, the embryos were washed and equilibrated in NTMT buffer followed by coloration with 4-nitro blue tetrazolium (NBT, Roche) and 5-bromo-4chloro-3-indolyl-phosphate, 4-toluidine salt (BCIP, Roche) in NTMT buffer. A stop solution (PBS pH 5.5, 1mM EDTA) was used to end the coloration reaction and embryos were placed in 40% glycerol for imaging on a dissecting microscope (Digital Sight Ds-Fi2, Nikon instruments). Six embryos were analyzed per time point for each probe.

### Immunohistochemistry

Immunohistochemistry was performed as previously described.^55^ Primary antibodies used in this paper are described in Table 2. Alexa Fluor 488 goat anti-mouse, 488 goat anti-rabbit, 546 goat anti-rabbit, and 546 goat anti-mouse secondary antibodies (Molecular Probes, Invitrogen) were all used at 1:200 dilution. Nuclei were visualized by counterstaining with DAPI (1:10,000 dilution). Samples were mounted in 60% glycerol in PBS. Images were taken at 20x and 40x on an inverted fluorescent microscope (Eclipse Ti-U, Nikon instruments). At least six sections were analyzed on each slide and for each antibody.

**Table 2:** Antibodies used for immunohistochemistry (IHC).

| Antibody | Labels | Raised in | Dilution | Acquired from |
| --- | --- | --- | --- | --- |
| Zpr-1 | red/green cones | mouse | 1:20 | ZIRC |
| 4C12 | rods | mouse | 1:100 | James Fadool<br>FSU, Tallahassee, FL |
| Zrf-1 | Müller glia | mouse | 1:5000 | ZIRC |
| N1C1 | Capn5 | rabbit | 1:100 | GeneTex |
| GFP | GFP | rabbit | 1:1000 | abcam |

### Light damage experiment

Acute light damage (LD) was performed essentially as previously described.^37,53^ Briefly, 18 month old, albino zebrafish were dark adapted for 14 days. Fish were then placed in a 2.8 liter clear plastic tank surrounded by four 250W halogen bulbs placed 7 inches away, which collectively produced 20,000 lux of light. A bubbler, cooling fan, and water circulatory system were used to maintain the water level and keep the temperature under 32°C. The fish were maintained in constant light for 3 days, at which point they were returned to normal lighting conditions. Fish were collected at various time points during and after light damage (LD), and the left eyes were dissected for cryosectioning and in situ hybridization, whereas the right eyes were dissected and processed for RNA extraction followed by RT-PCR and qPCR. The fish were collected at 3 days LD, and at 2 and 7 days post LD. The light damage experiment was repeated three times, and three fish were collected for each time point.

### Quantification and Statistical analysis

For comparisons between groups, statistical significance was determined using the Student’s t-test with p<0.05 considered as significant. For all graphs, data are represented as the mean +/-the standard deviation (s.d.).

## Acknowledgments

We are grateful to Charles Mashburn, Vimala Bondada, and James Geddes at the University of Kentucky Spinal Cord and Brain Injury Research Center for technical assistance and helpful discussions. The authors would also like to thank Sara Perkins and Chris Mitchell for zebrafish care, and Kayla Titialii for editorial assistance.

## Funding

This work was supported by a grant from the National Institutes of Health (R01EY021769, A.C.M.) and the University of Kentucky Lyman T. Johnson fellowship (C.E.C).

